# Drug repurposing screen identifies masitinib as a 3CLpro inhibitor that blocks replication of SARS-CoV-2 *in vitro*

**DOI:** 10.1101/2020.08.31.274639

**Authors:** Nir Drayman, Krysten A. Jones, Saara-Anne Azizi, Heather M. Froggatt, Kemin Tan, Natalia Ivanovna Maltseva, Siquan Chen, Vlad Nicolaescu, Steve Dvorkin, Kevin Furlong, Rahul S. Kathayat, Mason R. Firpo, Vincent Mastrodomenico, Emily A. Bruce, Madaline M. Schmidt, Robert Jedrzejczak, Miguel Á. Muñoz-Alía, Brooke Schuster, Vishnu Nair, Jason W. Botten, Christopher B. Brooke, Susan C. Baker, Bryan C. Mounce, Nicholas S. Heaton, Bryan C. Dickinson, Andrzej Jaochimiak, Glenn Randall, Savaş Tay

**Affiliations:** The Pritzker School for Molecular Engineering, The University of Chicago, Chicago, IL, USA; The Department of Chemistry, The University of Chicago, Chicago, IL, USA; The Department of Molecular Genetics and Microbiology, Duke University, Durham, NC, USA; The Department of Biochemistry and Molecular Biology, The University of Chicago, Chicago, IL, USA; The Cellular Screening Center, The University of Chicago, Chicago, IL, USA; Department of Microbiology, Ricketts Laboratory, University of Chicago, Chicago, IL, USA; Center for Structural Genomics of Infectious Diseases, Consortium for Advanced Science and Engineering, University of Chicago, Chicago, IL 60637, USA; Structural Biology Center, X-ray Science Division, Argonne National Laboratory, Argonne, IL 60439, USA; Department of Medicine, Division of Immunobiology, Larner College of Medicine, University of Vermont, Burlington VT, 05405, USA; Department of Microbiology and Molecular Genetics, Larner College of Medicine, University of Vermont, Burlington VT, 05405, USA; Vaccine Testing Center, Larner College of Medicine, University of Vermont, Burlington, VT, 05405 USA; Department of Molecular Medicine, Mayo Clinic, Rochester, MN 55905 USA; Department of Microbiology, University of Illinois at Urbana-Champaign, Urbana, Illinois, USA; Carl R. Woese Institute for Genomic Biology, University of Illinois at Urbana-Champaign, Urbana, Illinois, USA; Department of Microbiology and Immunology, Stritch School of Medicine, Loyola University Chicago, Maywood, Illinois 60153, USA

## Abstract

There is an urgent need for anti-viral agents that treat SARS-CoV-2 infection. The shortest path to clinical use is repurposing of drugs that have an established safety profile in humans. Here, we first screened a library of 1,900 clinically safe drugs for inhibiting replication of OC43, a human beta-coronavirus that causes the common-cold and is a relative of SARS-CoV-2, and identified 108 effective drugs. We further evaluated the top 26 hits and determined their ability to inhibit SARS-CoV-2, as well as other pathogenic RNA viruses. 20 of the 26 drugs significantly inhibited SARS-CoV-2 replication in human lung cells (A549 epithelial cell line), with EC50 values ranging from 0.1 to 8 micromolar. We investigated the mechanism of action for these and found that masitinib, a drug originally developed as a tyrosine-kinase inhibitor for cancer treatment, strongly inhibited the activity of the SARS-CoV-2 main protease 3CLpro. X-ray crystallography revealed that masitinib directly binds to the active site of 3CLpro, thereby blocking its enzymatic activity. Mastinib also inhibited the related viral protease of picornaviruses and blocked picornaviruses replication. Thus, our results show that masitinib has broad anti-viral activity against two distinct beta-coronaviruses and multiple picornaviruses that cause human disease and is a strong candidate for clinical trials to treat SARS-CoV-2 infection.

## Introduction

On January 2020, SARS-CoV-2 was identified as the causative agent of a new respiratory syndrome that was later named Corona Virus Disease 19 (COVID-19)^1^. The virus has rapidly spread throughout the world, causing an ongoing pandemic, with over 25 million confirmed cases and over 850,000 deaths (as of August 30^th^ 2020)^2^. SARS-CoV-2 is a member of *Coronaviridae*, a family of enveloped, single-strand, positive-sense RNA viruses with a relatively large genome^3^. This family includes both human and animal pathogens, including two other emerging human pathogens (SARS-CoV and MERS-CoV) as well four endemic human viruses that are the second most common cause of the common cold (HCoV-OC43, 229E, NL63 and HKU1)^4^.

Upon entry into the host cell cytoplasm, the viral genome is translated into a polyprotein that must be cleaved into the individual viral proteins for infection to proceed. This cleavage is mediated by two virally encoded proteases: the main viral protease, known as Mpro, 3CLpro or non-structural protein 5 (nsp5) and a second protease known as the papain-like protease, PLpro, a domain within nsp3^3^. There is much interest in developing *de-novo* inhibitors to target these proteases^5–10^ but the full clinical development of such drugs is a time-consuming process, usually requiring several years.

While several promising vaccine candidates have entered human clinical trials^11^, their success is not guaranteed. Moreover, even if some are approved, it will be some time until both production and vaccination efforts can reach the levels needed to control this global outbreak. Therefore, there is a need for new treatment options for COVID-19. While the antiviral remdesivir, an RNA-dependent RNA-polymerase inhibitor originally developed to treat Hepatitis C virus and later repurposed for Ebola, did receive an emergency use authorization from the FDA due to its ability to shorten COVID-19 hospitalization times^12^, there are currently no FDA-approved specific antivirals for treating SARS-CoV-2 infection and the introduction of additional antivirals would help to reduce morbidity and mortality.

One obvious way to speed up drug discovery is through the use of drug-repurposing screens, looking to identify safe-in-human drugs with potential anti-coronavirus properties^9,13,14^. Here, we employed a three-tiered approach for drug repurposing, to identify potential broad-spectrum anti-coronavirus drugs that specifically target the viral main protease, 3CLpro. We first screened a library of 1,900 safe-in-human drugs for their ability to inhibit HCoV-OC43 infection. HCoV-OC43 is a biosafety level 2 (BSL2) agent and a relatively safe model organism that enables the study of coronavirus infections outside of high biocontainment facilities. We then validated the anti-viral properties of the top hits against SARS-CoV-2, and finally investigated the drugs’ ability to inhibit the enzymatic activity of the main SARS-CoV-2 protease 3CLpro.

We report here on the identification of 20 safe-in-human drugs that are able to inhibit SARS-CoV-2 infection in human lung cells and on the mechanism of action of one of these, masitinib (a tyrosine-kinase inhibitor) - which we find to be a bone-fide 3CLpro inhibitor with broad anti coronaviruses and picornaviruses activity.

## Results

### A drug repurposing screen against the human beta coronavirus OC43 identifies multiple drugs that are effective against SARS-CoV-2

We began by screening a library of 1,900 clinically used drugs, either approved for human use or with extensive safety data in humans (Phase 2 or 3 clinical trials), for their ability to inhibit OC43 infection of the human lung epithelial cell line A549 (expressing an H2B-mRuby nuclear reporter). One day after plating, cells were infected at an MOI of 0.3, incubated at 33^°^C for 1 hour and drugs were added to a final concentration of 10μM. Cells were then incubated at 33^°^C for 4 days, fixed and stained for the presence of the viral nucleoprotein (Fig. 1A). We imaged the cells at day 0 (following drug addition) and day 4 (after staining) to determine the drugs effect on cell growth and OC43 infection.

**Figure 1.**
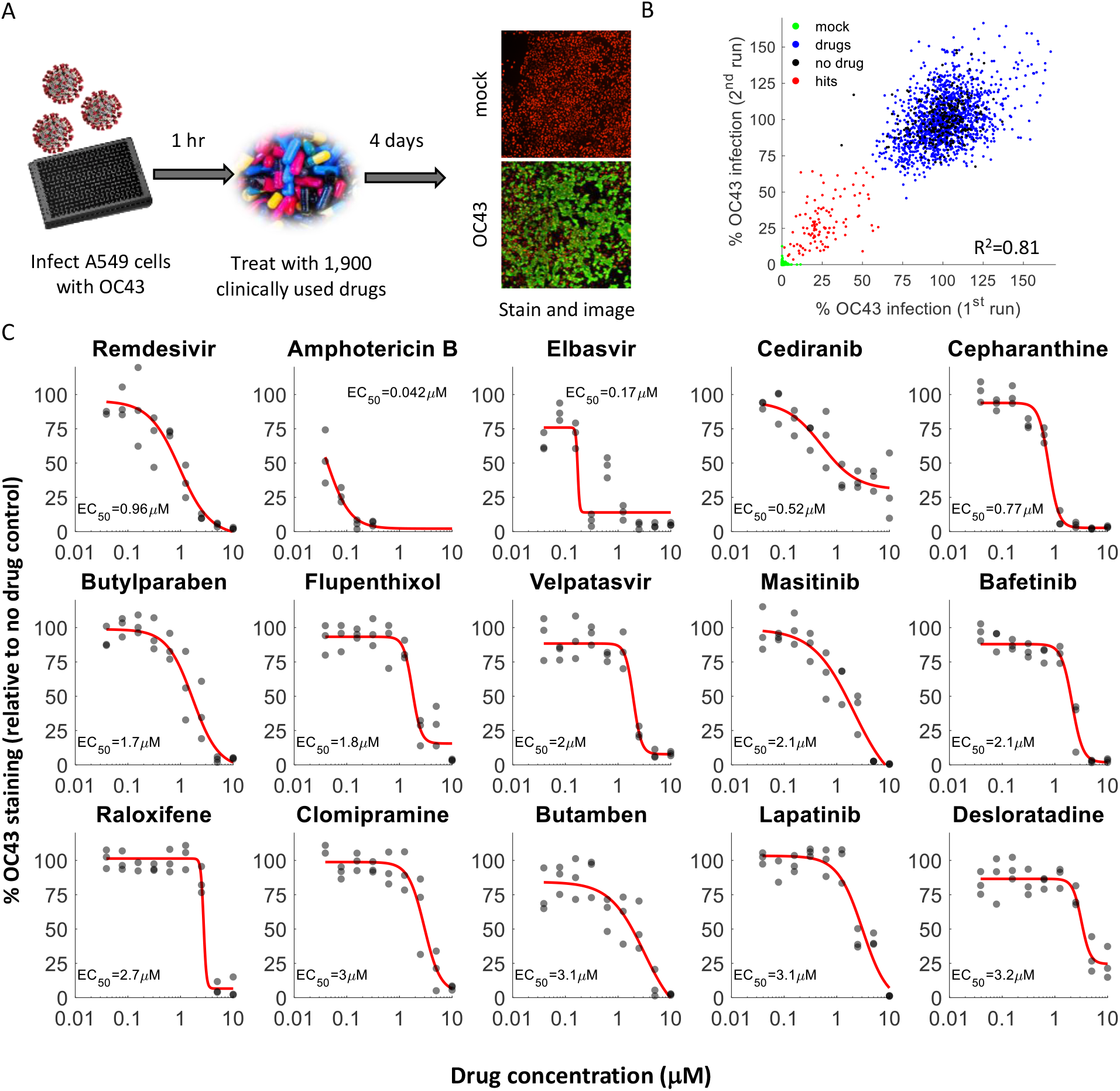
A drug repurposing screen identifies multiple safe-in-human drugs that inhibit OC43 infection. **A.** Schematic of the screen. A549 cells expressing H2B-mRuby were infected with OC43 (MOI 0.3), treated with drugs, incubated for 4 days at 33°C, and stained for the viral nucleoprotein. **B.** Screen results showing the %OC43 staining of mock-infected cells (green), no-drug controls (black), drugs with no effect on OC43 infection (blue), and screen hits (red). Overall agreement between the two repeats is high (R^2^=0.81) **C.** Dose response curves of remdesivir and the top hits from the screen, n = 3. Individual measurements are shown as semi-transparent circles (note that some circles overlap).

We repeated the screen twice and identified 108 drugs that significantly reduced OC43 infection without significant cellular toxicity (Fig. 1B, Fig. S1, Table S1). Of our top hits, we selected 29 drugs for further validation. We determined the EC50 values (drug concentration required to reduce infection by 50%) of these drugs against OC43 infection (Fig. 1C, Fig. S2). All of the drugs except erythromycin inhibited OC43 infection in a dose-dependent manner, with EC50 values ranging from 0.04-7μM. The most potent drugs in inhibiting OC43 infection were amphotericin B (an anti-fungal drug, EC50=0.04μM), elbasvir (an anti-viral against HCV, EC50=0.17μM), cediranib (an EGFR inhibitor, EC50=0.52μM) and cepharanthine (an anti-inflammatory drug, EC50=0.77μM). Remdesivir inhibited OC43 infection in the same cells with an EC50 value of 0.9μM.

We proceeded to determine the EC50 values for 26 of these drugs against SARS-CoV-2 infection. In a high biocontainment (BSL3) facility, A549 cells over-expressing the angiotensin-converting enzyme 2 (ACE2) receptor were treated with the drugs for 2 hours, infected with SARS-CoV-2 at an MOI of 0.5, incubated for 2 days, fixed, and stained for the viral spike protein (as a marker of SARS-CoV-2 infection). After staining, the cells were imaged under a microscope to quantify the fraction of infected cells. Of the 26 drugs tested, 20 (77%) inhibited SARS-CoV-2 infection in a dose dependent manner (Fig. 2A, Fig. S3). Interestingly, the most potent drugs against OC43 infection (amphotericin B and elbavir) did not inhibit SARS-CoV-2 infection. The pleiotropic drug cepharanthine was the most potent inhibitor of SARS-CoV-2 infection (EC50=0.13μM), followed by flupenthixol (an anti-psychotic, EC50=0.56μM) and desloratidine (an anti-histamine, EC50=0.9μM). In comparison, remdesivir inhibited SARS-CoV-2 infection with an EC50 value of 0.1μM. A comparison of the EC50 values obtained against OC43 and SARS-CoV-2, as well as the chemical structures of the drugs, is shown in Table S2. Many of the identified drugs are anti-psychotic and anti-allergic drugs that share a similar tricyclic structure, in agreement with the previous identification of olanzapine (a tricyclic anti-depressant) as an anti-SARS-CoV-2 drug^13^.

**Figure 2.**
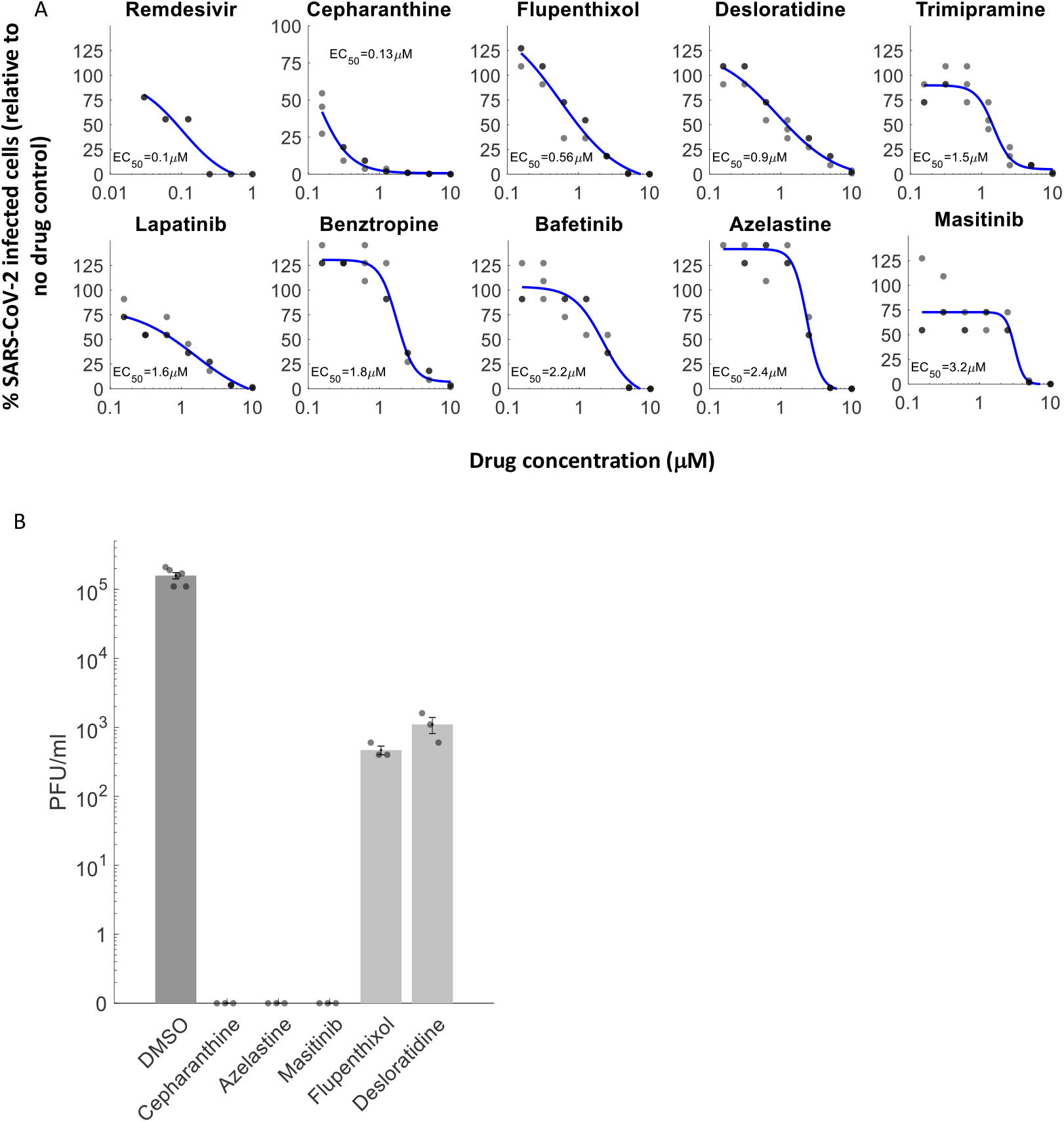
Discovery of repurposed drugs that inhibit SARS-CoV-2 replication in human lung cells. Of the 26 drugs that inhibited OC43 and tested against SARS-CoV-2, 20 inhibited SARS-CoV-2 replication in a dose-dependent manner, showing good concordance between OC43 and SARS-CoV-2 inhibition. **A.** A549 cells over-expressing ACE2 were pre-treated with indicated drugs for 2 hours, infected with SARS-CoV-2 (MOI 0.5) and incubated for 2 days. Cells were stained for the presence of the spike protein and the % of infected cells was analyzed. Most of the drugs effective against OC43 showed similar effectivity against SARS-CoV-2, n=3. Individual measurements are shown as semi-transparent circles (note that some circles overlap). **B.** Effect of selected drugs on SARS-CoV-2 progeny production. Cells were treated with 10 μM of the indicated drugs for 2 hours, infected with SARS-CoV-2 (MOI 0.5) and cell supernatant were collected for titration 2 days later, n = 3. Individual measurements are shown as semi-transparent circles. All drugs showed a statistically significant (p-values<0.001, one-tailed t-test, FDR-corrected) reduction in viral titers.

We further evaluated the effect of several drugs on the production of viable progeny viruses. Cells were treated with 10μM of the drugs for 2 hours, infected at an MOI of 0.5 and the supernatant collected 2 days after for titration (Fig. 2B). Cepharanthine, azelastine, and masitinib completely eliminated SARS-CoV-2 progeny production (>5-logs decrease), while flupenthixol and desloratidine reduced viral titers by about 2-logs. Ultimately, our screen identified 20 safe-in-human drugs that are able to inhibit both OC43 and SARS-CoV-2 infection *in vitro*.

### Masitinib is a bone-fide 3CLpro inhibitor

We next examined the drugs ability to inhibit the SARS-CoV-2 main protease (also known as 3CLpro, Mpro and nsp5). 3CLpro is an attractive target for anti-viral drugs, as it indispensable for the viral replication cycle and is well conserved among coronaviruses^15^. We first tested the ability of 20 drugs that inhibited both SARS-CoV-2 and OC43 infection to inhibit 3CLpro activity in 293T cells transfected with a FlipGFP reporter system^16^ at a single concentration of 10μM. In this assay, 3CLpro cleavage of the FlipGFP reporter is needed to produce GFP fluorescence, and thus the level of GFP+ cells reports on 3CLpro activity. 8 drugs showed a statistically significant decrease in the percentage of GFP-expressing cells - retapamulin, conivaptan, bafetinib, raloxifene, imatinib, lapatinib, vilazodone and masitinib (Fig. 3A, Fig. S4). Standing out among these was masitinib, a c-kit/PDGFR inhibitor in multiple phase 3 clinical trials, which completely inhibited 3CLpro activity. Masitinib preferentially binds and inhibits the mutant form of c-kit^17^ and has been approved for treatment of mast-cell tumors in dogs^18^. Masitinib was also evaluated in phase 2 and 3 clinical trials in humans for the treatment of cancer^19^, asthma^20^, Alzheimer’s^21^, multiple sclerosis^22^ and amyotrophic lateral sclerosis^23^.

**Figure 3.**
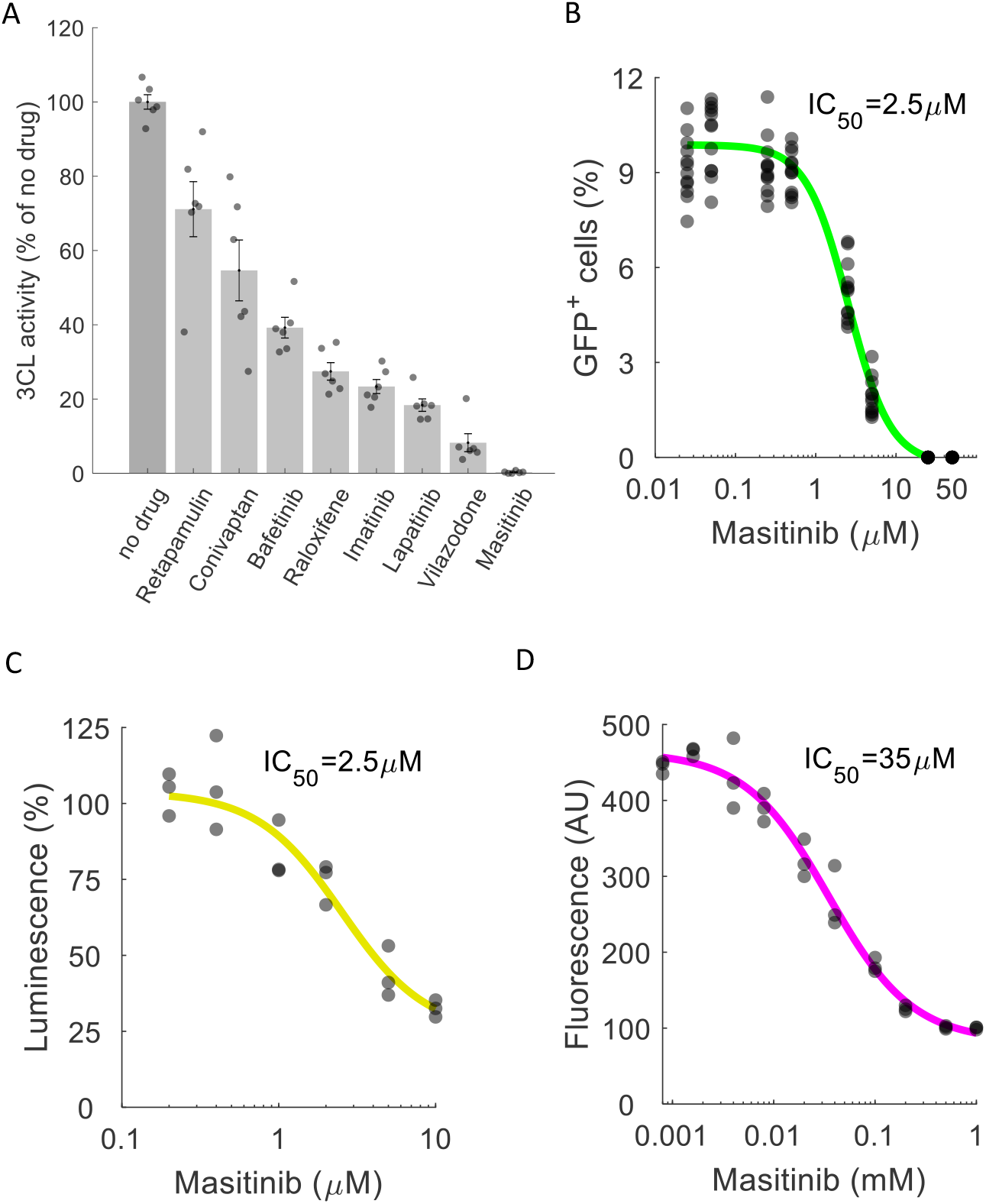
Masitinib inhibits SARS-CoV-2 3CLpro enzymatic activity. **A.**A FlipGFP reporter assay was performed to screen for potential inhibition of 3CLpro by the identified drugs at a single concentration (10μM). Shown are the drugs that showed a statistically significant reduction in 3CLpro activity (p-value<0.05, one-tailed t-test, FDR-corrected). n=6. The data for the remaining tested drugs is shown in Figure S4. Individual measurements are shown in circles. Bars depict mean ± s.e. Masitinib treatment completely inhibited 3CLpro activity. **B.** Dose-response curve for 3CLpro inhibition by masitinib using the FlipGFP reporter assay, n = 6. Individual measurement shown as circles. **C.** Dose-response curve for 3CLpro inhibition by masitinib using a luciferase reporter assay, n = 3. Induvial measurement shown as circles **D.** Dose-response curve for 3CLpro inhibition in a cell-free assay using purified 3CLpro and a flurogenic peptidic substrate, n = 3. Individual measurement shown as circles.

Therefore, we proceeded to determine the IC50 value (the drug concentration that causes a 50% reduction in enzymatic activity) of masitinib inhibition of 3CLpro activity in two distinct cellular assays; the same FlipGFP reporter assay described above (Fig. 3B), as well as a luciferase reporter assay adapted for SARS-CoV-2^24^ (Fig. 3C). These assays, performed independently at the University of Chicago and Duke University, determined the IC50 value to be 2.5μM (Fig. 3B,C), similar to the EC50 values determined against OC43 (2.1μM, Fig. 1C) and SARS-CoV-2 (3.2μM, Fig. 2A) infections, suggesting that masitinib inhibition of coronavirus infection is achieved by inhibiting 3CLpro activity.

We additionally evaluated the IC50 value of masitinib using a novel cell-free assay with purified 3CLpro and a methyl-amino coumarin (AMC)-tagged peptide, whose fluorescent signal is uncaged upon 3CLpro proteolytic activity (Fig. 3D, IC50 = 35 μM. See Methods for complete description of the assay). While this assay determined a higher IC50 value than the cell-based assays, it strongly suggested that direct interaction of masitinib with the viral protease is responsible for its effects on SARS-CoV-2 replication.

### Masitinib inhibits 3CLpro by directly binding to its active site

To obtain further mechanistic understating of the mode of 3CLpro inhibition by masitinib we determined the high-resolution structure of masitinib-bound 3CLpro using X-ray crystallography (Fig. 4). The structure indicates that masitinib binds non-covalently into the shallow, elongated grove between domains I and II of 3CLpro and is crossing the 3CLpro active site. The enzyme is a dimer and both active sites are occupied by masitinib. Specifically, masitinib’s pyridine ring packs into the S1 substrate pocket of 3CLpro peptide recognition site^25^. In addition to hydrophobic and Van der Waals interactions between the ring and its surrounding pocket-forming residues, it forms a hydrogen bond with His163, located at the bottom of the S1 pocket. Masitinib’s aminothiazole ring forms a hydrogen bond with the carbonyl of His164 and interacts directly with the key catalytic residue - Cys145. The second catalytic residue, His41, forms a nearly perfect π-π stacking with masitinib’s hydrophobic toluene ring that occupies the S2 binding pocket. These three active groups (pyridine, aminothiazole and toluene rings) contribute the majority of interactions between masitinib and 3CLpro, bind the key active site residues and effectively block the peptide substrate access to the protease catalytic dyad^26^, thus preventing polyprotein cleavage.

**Figure 4.**
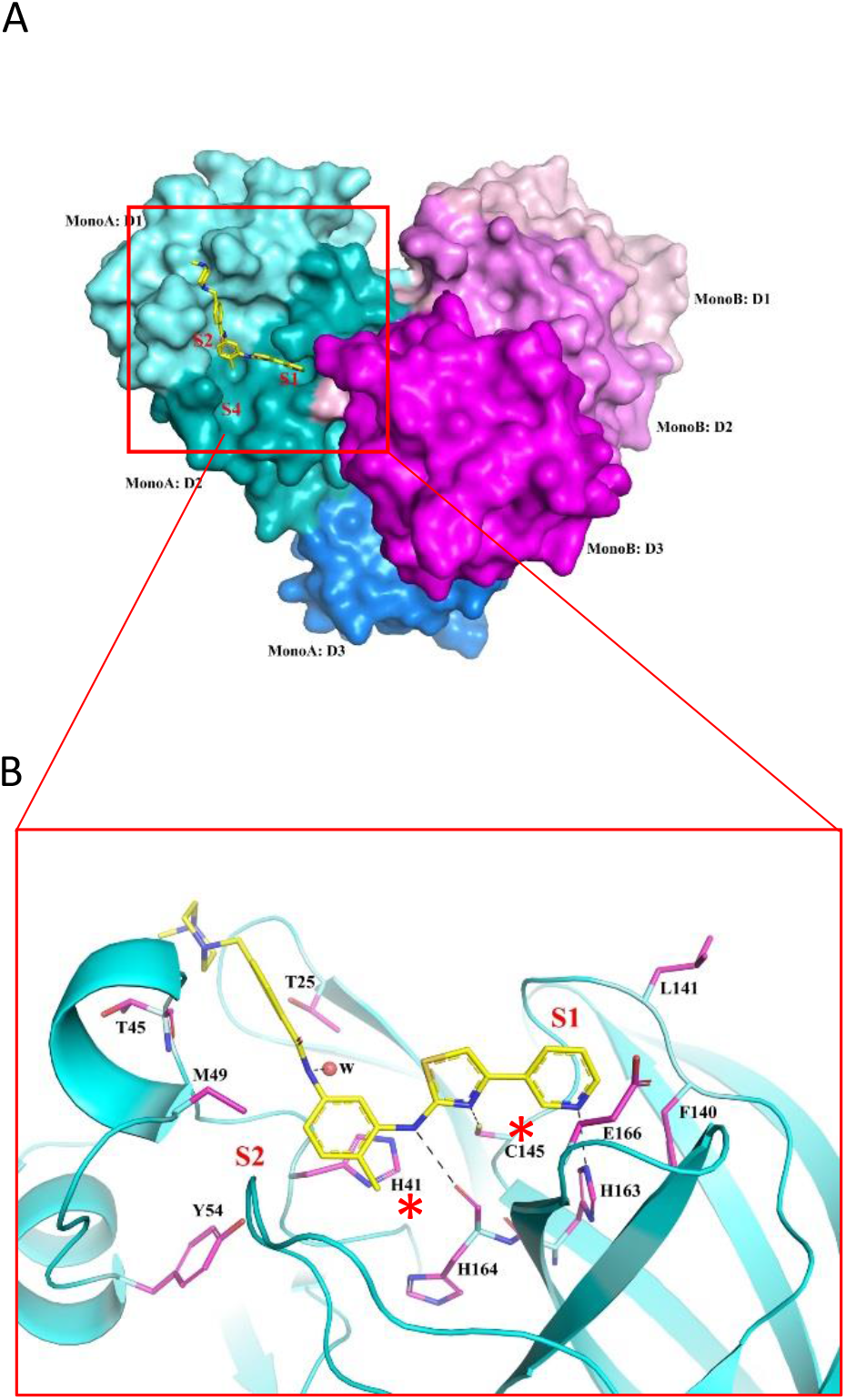
Binding mode and inhibition of 3CLpro by masitinib. The dimer formation, domain structure, and masitinib binding site of SARS-CoV-2 3CLpro. Domains I, II and III (D1-D3) of the monomer A of a 3CLpro dimer are colored in cyan, teal and light blue, respectively. The corresponding three domains of monomer B are colored in light pink, magenta and purple, respectively. In monomer A, the inhibitor masitinib is drawn in stick format, bound to the active site between D1 and D2. The sites of three binding pockets S1, S2, and S4 are marked in red. **B.** The interaction of masitinib with 3CLpro. The ribbon diagram shows details of some interactions formed between masitinib and 3CLpro at the active site. Masitinib is drawn in stick format with its C atoms colored in yellow. Key pocket forming or interacting residues of 3CLpro are also presented in stick format with their C atoms colored in purple. Hydrogen bonds are drawn in black dashed lines. The sites of binding pockets S1 and S2 are marked in red. The two catalytic residues are marked by red asterisks.

Taken together, our results show that masitinib, originally designed as a tyrosine-kinase inhibitor and considered for treatment of a number of human diseases, harbors potent anti-coronavirus activity through its direct binding to and inhibition of the virus main protease.

### Masitinib blocks the replication of picornaviruses through the inhibition of their 3C protease

Since masitinib directly binds and inhibits the coronaviruses 3CL protease, we asked if it is also effective against the 3C protease of picornaviruses (human pathogens that cause a range of diseases including meningitis, hepatitis, and poliomyelitis) given the extensive structural homology and substrate specificity shared between these viral proteases^27^. Using a luciferase reporter assay^28^, we indeed found that masitinib significantly inhibited the activity of the 3C protease in cells (Fig. 5A). Masitinib was also effective in blocking the replication of multiple picornaviruses (Fig. 5B) but not of other RNA viruses (Fig. S5). Thus, we conclude that masitinib is a relatively broad-spectrum anti-viral, able to inhibit multiple corona- and picorna-viruses, but not other RNA viruses that do not rely on a 3CL-like protease to complete their life cycle.

**Figure 5.**
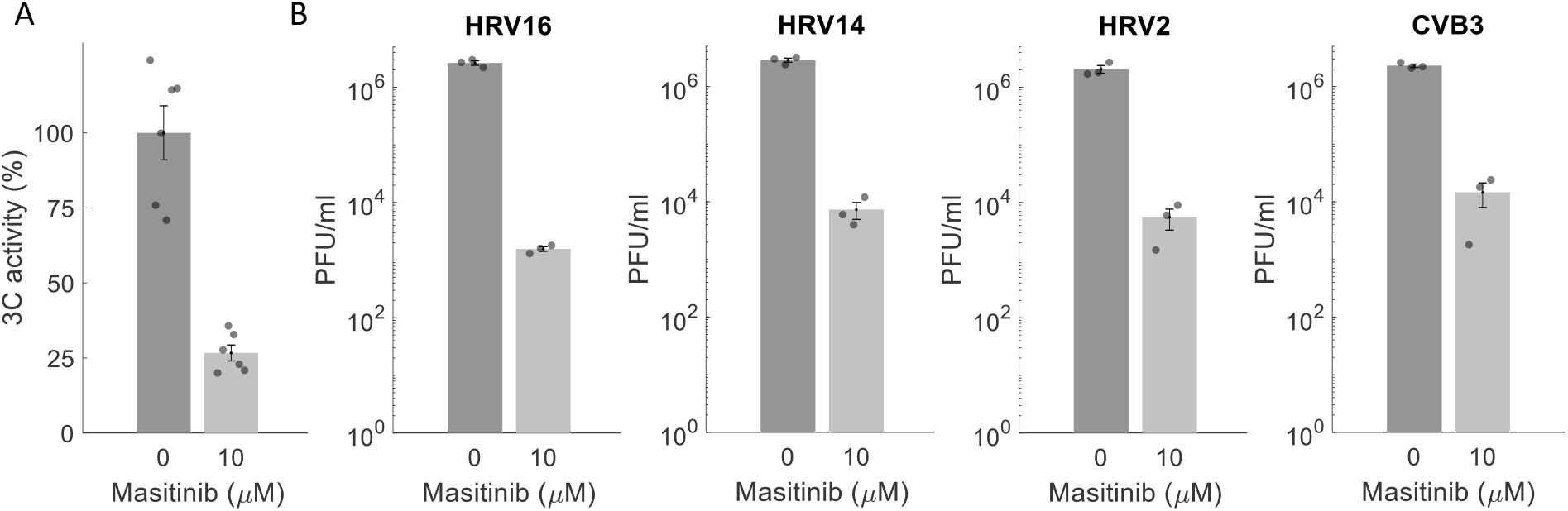
Masitinib inhibits picornaviruses through inhibition of their 3C protease. **A.** A luciferase reporter assay was performed to investigate masitinib ability to inhibit the proteolytic activity of picornaviruses 3C (derived from CVB3). n=6, p-value=7X10^−6^ (one-tailed t-test). **B.** Huh7 cells were treated with 10μM masitinib for 2 hours, infected with coxsackievirus B3 (CVB3) or human rhinoviruses 2, 14 and 16 (HRV 2,14, 16) at an MOI of 0.01 and the supernatant collected for titration 24 hours later. n=3, p-values< 0.001 (one-tailed t-test, FDR-corrected).

## Discussion

In conclusion, we have shown that OC43, a BSL-2 pathogen that can be readily studied in most virological labs, is a good model system to screen potential anti-viral drugs against SARS-CoV-2 infection, since most drugs that inhibited OC43 replication also inhibited SARS-CoV-2 in our measurements. We identified 20 safe-in-human drugs that should be considered for advancement for clinical trials and identified masitinib as a direct anti-viral agent that binds and inhibits 3CLpro.

While masitinib binds to 3CLpro in a non-covalent manner, it shows better efficacy against SARS-CoV-2 replication *in vitro* than the recently reported covalent, pre-clinical, 3CLpro inhibitor 13b^29^. While masitinib’s EC50 value is higher than that of two other, pre-clinical, covalent inhibitors, 11a and 11b^10^, it showed superior inhibition of progeny production at 10μM (over 5-logs for masitinib, compared to 2-logs for 11a and 11b).

It is important to note that in addition to the direct anti-viral effect we have uncovered here, masitinib has been shown to decrease airway inflammation and improve lung functions in a feline model of asthma^30^. Given that a main pathology of SARS-CoV-2 is ARDS (acute respiratory distress syndrome), the combined anti-viral and anti-inflammatory properties of masitinib might prove highly beneficial for treating COVID-19.

Though here we highlighted masitinib, given its direct inhibitory effect on the viral main protease, we emphasize that the other drugs we identified as SARS-CoV-2 inhibitors should also be investigated more. Topical drugs in particular represent an interesting class of drugs for further exploration. These drugs could be considered for use as prophylactics in high risk populations, such as healthcare workers, due to their ease of use, favorable safety profile, and local delivery of high doses to areas of viral replication. In addition to azelastine (a nasally administered antihistamine used to treat rhinitis), which we have shown inhibits both OC43 (Fig. 1C) and SARS-CoV-2 (Fig. 2), several of the identified hits in our primary screen against OC43 are also administered topically to either the nose or lungs (Table S1). These include salmeterol, budesonide, beclomethasone, fluticasone, and xylometazoline.

Future efforts should evaluate the efficacy of the identified drugs in preventing or treating COVID19 patients as well as decipher the mechanism by which these drugs are able to inhibit beta-coronaviruses replication. While any single agent anti-viral therapy is likely to cause viral adaption and the emergence of resistance mutants, a previous work suggested that CoVs that escape 3CLpro inhibition by mutating the protease show decreased replication and pathogensis^31^. The identification of direct anti-virals that target multiple distinct aspects of the viral life cycle (such as the polymerase, proteases, helicase etc.) will allow the design of combination therapies that have proven highly beneficial against other RNA viruses, such as HIV and HCV.

**TABLE 1.** Summary of crystallographic data

| **Data collection statistics** | **M<sup>pro</sup> + masitinib** |
| --- | --- |
| Space group | C2 |
| Unit cell (Å, °) | $a = 98.55$, $b = 81.23$,<br>$c = 51.85$, $\beta = 114.6$ |
| MW Da (residue) | 33,796 (306) |
| Mol (AU) | 1 |
| Wavelength (Å) | 0.9791 |
| Resolution | 1.60 |
| Number of unique reflections | 48,121 |
| R<sub>merge</sub> (%) | 8.4 (87.7)<sup>1</sup> |
| Completeness (%) | 98.5 (95.4)<sup>1</sup> |
| Redundancy | 3.5 (2.3)<sup>1</sup> |
| I/($\sigma$) | 26.3 (1.1)<sup>1</sup> |
| Solvent content (%) | 56.0 |
| **Phasing** |  |
| Resolution range (Å) | 43.5 – 3.00 |
| Correlation coefficient<sup>2</sup> (%) | 0.58 |
| **Refinement** |  |
| Resolution range (Å) | 43.5 – 1.60 |
| Number of reflections | 47,567 |
| Completeness (%) | 97.1 |
| R<sub>work</sub>/R<sub>free</sub> (%) | 16.8/19.2 |
| No. of Atoms | 2,363/249 |
| Bond lengths (Å) | 0.016 |
| Bond angles (deg) | 1.431 |
| B-factors (Å<sup>2</sup>) (main/side chain) | 32.73/52.9 |
| Wilson B-factor (Å<sup>2</sup>) | 23.98 |
| **Molprobity validation** |  |
| Ramachandran outliers (%) | 0.33 |
| Ramachandran Favored (%) | 98.34 |
| Rotamer outliers (%) | 0.77 |
| Clashscore | 2.72 |
| MolProbity score | 1.06 |
| **PDB ID** | 7JU7 |

<sup>1</sup> Last resolution bin (1.60-1.63 Å); <sup>2</sup> Molecular replacement method.

## Supporting information

Table_S2_EC50_OC43_SARS_CoV_2

Table_S1_screen_results

## Acknowledgments

ND is a recipient of the Human Frontiers Science Program (HFSP) post-doctoral fellowship. We thank the members of the SBC at Argonne National Laboratory, especially Darren Sherrell and Alex Lavens for their help with setting beamline and data collection at beamline 19-ID. We would like to thank Lucas F. Welk for assistance in purification of 3CLpro. Funding for this project was provided in part by federal funds from the National Institute of Allergy and Infectious Diseases, National Institutes of Health, Department of Health and Human Services, under Contract HHSN272201700060C (to AJ) and by the DOE Office of Science through the National Virtual Biotechnology Laboratory, a consortium of DOE national laboratories focused on response to COVID-19, with funding provided by the Coronavirus CARES Act (to AJ). EAB acknowledges funding by NIH P20GM125498 (UVM Translational Global Infectious Disease Research Center) and NIH P30GM118228-04 (UVM Center for Immunology and Infectious Disease). JWB acknowledges funding from the Office of the Vice President for Research at the University of Vermont and NIH grants R41AI132047, R21AI154198, and U01AI1141997.The use of SBC beamlines at the Advanced Photon Source is supported by the U.S. Department of Energy (DOE) Office of Science and operated for the DOE Office of Science by Argonne National Laboratory under Contract No. DE-AC02-06CH11357. The rLCMV Armstrong 53b reverse genetics system was generously provided by Juan Carlos de la Torre.

## Author contributions

ND conducted the OC43 drug screening and dose-response experiments, analyzed data, generated figures, wrote the manuscript and supervised the project. SAA synthesized the 3CL fluorogenic substrate with assistance from RSK. KAJ conducted and analyzed the 3CLpro kinetic assay and 3CL luciferase assay. HMF conducted and analyzed the 3CL FlipGFP assay. KT and NI conducted and analyzed the x-ray crystallography and generated figures. SC conducted the OC43 drug screening and dose-response experiments. VN, SD and KF conducted and analyzed the SARS-CoV-2 dose response and tittering experiments. MRF, VM and BCM conducted and analyzed the picornaviruses experiments. EAB, MMS and JWB conducted and analyzed the LCMV experiments. RJ designed the recombinant 3CLpro purification system. MMA conducted and analyzed the Measles experiments. BS and VN assisted in data analysis. CBB conducted and analyzed the IAV experiments. SCB designed the 3CLpro luciferase assay. NSH supervised the 3CL FLipGFP assay and wrote the manuscript. BCD supervised the 3CL kinetics assay and luciferase assay and wrote the manuscript. AJ supervised the x-ray crystallography and wrote the manuscript. GR supervised the SARS-CoV-2 experiments and wrote the manuscript. ST supervised the project and wrote the manuscript.

## Methods

### Cells

A549 expressing H2B-mRuby were generated by first infecting A549 cells (ATCC CCL-185) with a lentivirus (carrying H2B-mRuby), and FACS-sorting mRuby+ cells. They were maintained as a polyclonal population and grown in DMEM+10% BCS (Bovine Calf Serum). These cells were used for all OC43 infections. Ace2-A549^32^ cells were used for SARS-CoV-2 infections and were a kind gift of Benjamin TenOever, Mt Sinai Icahn School of Medicine. They were maintained in DMEM + 10% FBS. African green monkey kidney cells (Vero E6), kindly provided by J.L. Whitton, were maintained in DMEM supplemented with 10% FBS, 1% penicillin-streptomycin and 1% HEPES. We used Huh7 cells for picornaviruses infections. MDCK-SIAT1-TMPRSS2 cells, obtained from Jesse Bloom, were used for IAV infections. A549 cells maintained in 50:50 DMEM:F-12 media supplemented with 10% FBS and 1% penicillin-streptomycin were used for LCMV infections.

### Viruses

OC43 was obtained from ATCC (VR-1558) grown and titrated on A549-mRuby cells. SARS-CoV-2 (nCoV/Washington/1/2020) was provided by the National Biocontainment Laboratory, Galveston, TX. VeroE6 cells were used to propagate and titer SARS-CoV-2. CVB3 (Nancy strain), HRV 2, 14, and 16 were derived from full-length infectious clones and generated in Vero cells (NR-10385, BEI Resources, NIAID, NIH). Rhinovirus infectious clones were provided by Dr. Bill Jackson. Recombinant LCMV based on the Armstrong 53b strain was generated as previously described^33,34^. Working stocks were generated in Vero E6 cells, and the same cells were used to measure virus titers. The Measles Virus used was derived from the molecular cDNA clone of the Moraten/Schwartz vaccine strain^35^. The recombinant Measles virus was engineered to express firefly luciferase as previously described^36^.

### Drug screening

A549-mRuby cells were seeded (3,000 cells per well) in nine 384-well plates using Multidrop combi. Cells were seeded in a final volume of 30ul with DMEM+10% BCS. The following day, 20ul of OC43 were added (MOI 0.3) and incubated at 33°C, 5% CO^2^ for 1 hour. 50nl from the Selleck FDA-approved drug library (cat #L1300, Selleck) were added (1:1,000 dilution). Two columns (32 wells) were left uninfected and two columns were treated with DMSO and virus (no drug control). Cells were imaged using the IncuCyte S3 to measure cell numbers at day 0. Cells were incubated for 4 days at 33°C, 5% CO2 and were stained for OC43 nucleoprotein. All the following steps were performed at room temperature. Cells were fixed in 50ul 4% PFA/PBS for 15 min, blocked with 50ul 10% BSA+0.5% Triton X-100 in PBS for 30 minutes, stained with 50ul anti-OC43 (cat # MAB9013, Milipore) diluted 1:2,000 in 2% BSA+0.1% Triton X-100 in PBS for 1 hour, washed with 50ul PBS three times, stained with anti-mouse-AlexaFluor488 diluted 1:1,000 in 2% BSA+0.1% Triton X-100 in PBS for 1 hour, washed with 50ul PBS three times and imaged on the IncuCyte S3 (day 4). The screen was performed twice.

The following parameters were extracted from the images: number of cells at day 0, number of cells at ay 4 and total OC43 staining intensity at day 4. For analysis, OC43 staining intensity was normalized to the number of cells in the well and further normalized to the mean of the no drug controls, which was set to 100. We removed from analysis compounds that showed significant effect on cell growth. For each plate, we considered a drug as a putative hit if it reduced OC43 staining by over 3 standard deviations from the mean of the no drug controls. Drugs were considered hits if they were not toxic and reduced OC43 staining by over 3 standard deviations in both repeats. The full list of screened drugs, as well as their effect on cell growth and OC43 staining is summarized in Table S1.

### Dose-response analysis for OC43 and SARS-CoV-2 infection

Dose-response analysis of OC43 infection was done similarly to the drug screening, except cells were seeded at a concentration of 5,000 cells per well and the media contained 2% BCS instead of 10% BCS. OC43 staining was performed 2 days after infection and analyzed similarly to what was described for the drug screening. A sigmoid fit was used to extract EC50 values using Matlab. In a few cases we were unable to fit a sigmoid curve to the data and EC50 values were estimated from the graphs (these are denoted by ~ in Fig. S2).

All SARS-CoV-2 infections were performed in biosafety level 3 conditions at the Howard T. Ricketts Regional Biocontainment Laboratory. Ace2-A549 cells in DMEM +2% FBS were treated with drugs for 2 hours with 2-fold dilutions beginning at 10μM in triplicate for each assay. Cells were infected with an MOI of 0.5 in media containing the appropriate concentration of drugs. After 48 hours, the cells were fixed using 3.7% formalin, blocked and probed with mouse anti-Spike antibody (GTX632604, GeneTex) diluted 1:1,000 for 4 hours, rinsed and probed with anti-mouse-HRP for 1 hour, washed, then developed with DAB substrate 10 minutes. Spike positive cells (n>40) were quantified by light microscopy as blinded samples.

For SARS-CoV-2 plaque titers, cell supernatants from the infection described above were serially diluted (10-fold steps were used) and used to infect Vero E6 cells for 1 hour. Inoculum was removed and 1.25% methylcellulose DMEM solution was added to the cells and incubated for 3 days. Plates were fixed in 1:10 formalin for 1 hour, stained with crystal violet for 1 hour and were counted to determine plaque forming units (PFU)/ml.

### FlipGFP SARS-CoV-2 3CLpro Assay

293T cells were seeded 24 hours before transfection on poly-lysine treated plates. The next day, SARS-CoV-2 3CLpro plasmid, FlipGFP coronavirus reporter plasmid, Opti-MEM, and TransIT-LT (Mirus) were combined and incubated at room temperature for 20 minutes before being added to the cells. Drugs were applied to the cells at the indicated concentrations at the time of transfection. 24 hours after transfection, cells were fixed with 2% PFA at room temperature for 20 minutes and were incubated in 1:10,000 Hoescht 33342 (Life Technologies) in PBS at 4°C overnight. Quantification was performed by using the CellInsight CX5 (Thermo Scientific) equipment.

### 3CLpro luciferase reporter assay

Approximately 16 hours before transfection, 293T cells were plated in 96-well plates and grown to 70-80% confluency overnight. The next day, we transfected the cells with 37.5 ng pGlo-30F-VRLQS, 37.5 ng SARS-CoV2 3CLpro, and 2.5 ng pRL-TK (Promega) using Lipofectamine 2000 (Invitrogen) using manufacture’s recommendations. After 18 hours, masitinib (0-10μM) was added to the cells and incubated for an additional 6 hours before luciferase readout on a Biotek Synergy plate reader as previously described^24^. Briefly, 40 μL growth media was removed from every well and then 40 μL firefly assay buffer (Triton Lysis Buffer (50 mM Tris, pH 7.0, 75 mM NaCl, 3 mM MgCl_2_, 0.25% Triton X-100) containing 5 mM DTT, 0.2 mM coenzyme A, 0.15 mM ATP, and 1.4 mg/mL D-luciferin) was added to lyse the cells, and to provide the substrate for firefly luciferase. Firefly luminescence was read 10 minutes later and 40 μL Renilla assay buffer (45 mM EDTA, 30 mM sodium pyrophosphate, 1.4 M NaCl, 0.02 mM PTC124, 0.003 mM coelentrazine h (CTZ-h)) was added to stop firefly luciferase activity and provide the substrate for Renilla luciferase. Renilla luminescence was read 2-3 minutes after addition of the buffer. Firefly luciferase luminescence was normalized to the corresponding Renilla luciferase luminescence to generate normalized luminescence.

### 3CLpro kinetic assay

The cell-free inhibition assay was done in triplicates at 25°C using 96-well plates. Reactions containing the different concentrations of masitinib (0-1,000 μM) and 3CLpro enzyme (0.4 μM) in Tris-HCl pH 7.3, 1 mM EDTA were incubated for approximately five minutes. Reactions were then initiated with Ac-TVLQ-AMC probe substrate (40 μM), shaken linearly for 5 seconds. Fluorescence emission intensity (excitation λ: 364 nm; emission λ: 440 nm) was measured continuously (Synergy Neo2 Hybrid). Data were fit using a sigmoid curve fit in Matlab.

### Gene cloning, protein over-expression and purification

Cloning of 3CLpro from SARS CoV-2 was based upon the original cloning of SARS-CoV 3CLpro^37^. The gene coding for 3CLpro from SARS CoV-2 was cloned between an upstream MBP and a downstream sequence of GPHHHHHH. Detailed cloning of pCSGID-Mpro carrying 3CLpro from SARS CoV-2 is described in Kneller et al^26^.

pCSGID-Mpro was transformed into 100 ml of E. coli BL21(DE3)-Gold (Strategene) under selection of ampicillin (150mg/L) and grewn overnight at 37°C. The starter was then transferred to 4 L of LB-Miller, culture and was grown at 37°C with constant shaking (190 rpm). After reaching an OD_600_ of ~1, the shaker was set to 4°C. When temperature reached 18°C, IPTG and K_2_HPO_4_ was added to 0.2 mM and 40 mM respectively and the culture was marinated at 18°C. The cells were spun down at 4000g, resuspended in lysis buffer (500 mM NaCl, 5% (v/v) glycerol, 50 mM HEPES pH 8.0, 20 mM imidazole pH 8.0, 1mM TCEP) and kept frozen at −80°C.

Bacterial cells were lysed by sonication and debris were removed by centrifugation at 25,400 × *g* for 60min at 4^°^C. The clarified supernatant was mixed with 3 ml of Ni2^+^ Sepharose (GE Healthcare Life Sciences) equilibrated with lysis buffer. The suspension was applied to a Flex-Column (420400-2510) which was connected to a Vac-Man vacuum manifold. Unbound protein was washed out using controlled suction lysis buffer (160 ml). 3CLpro was eluted using 15 ml of buffer containing 500 mM NaCl, 5% (v/v) glycerol, 50 mM HEPES pH 8.0, 500 mM imidazole pH 8.0 and 1mM TCEP. The fractions containing 3CLpro were pooled, and rhinovirus 3C His_6_ tagged protease was added at a 1:25 protease:protein ratio and incubated at 4 °C overnight to cleave the C-terminal His_6_ tag, resulting in a 3CLpro with an authentic N and C-termini. 10 kDa MWCO filter (Amicon-Millipore) was used to concentrate the protein solution, which was subsequently applied to Superdex 75 column, pre-equilibrated with lysis buffer. The fractions containing 3CLpro were pooled together and run through 2 ml of Ni resin. We collected the flow through and replaced the lysis buffer with crystallization buffer (20 mM HEPES pH 7.5, 150 mM NaCl, 2 mM DTT (1,4-Dithiothreitol, Roche, Basel, Switzerland)) using a 10 kDa MWCO filter. 3CLpro solution was concentrated to 49 mg/ml, was aliquoted, frozen and stored at −80^°^C.

### Crystallization of Masitinib with SARS-CoV-2 3CLpro

Crystallization were carried out using previous protocols^38^. 3CLpro was mixed with 0.2M masitinib solution in DMSO. The final protein concentration was 6.25 mg/ml and inhibitor concentration was 8 times higher. This mixture was incubated for 1 hour (at room temperature) and spun down at 12,000 x *g* to remove precipitation. For crystallization, we utilized the sitting-drop vapor-diffusion method via a Mosquito liquid dispenser (TTP LabTech, Royston, UK) in 96-well CrystalQuick plates (Greiner Bio-One, Monroe, NC, USA) using a protein-to-matrix ratio of 1:1. ProPlex, PACT *premier* (Molecular dimensions, Cambridge, UK), and TOP96 (Anatrace, Maumee, OH, USA) screens were used for crystallization at 16°C. The first thin-plate crystals (obtained one day later in several conditions) were applied as seeding. The best crystals appeared in PACT B7 (0.2M Sodium Chloride, 0.1MES pH 6.0, 20% PEG 6000), TOP96 H8 (0.1M Ammonium Acetate, 0.1M Bis-Tris pH 5.5, 17% PEG 10000) and Top96 F11 (0.1M Bis-Tris pH6.5; 25% PEG 3350). Crystals selected for data collection were treated in their crystallization buffers supplemented with 10-18% glycerol and were subsequently flash-cooled in liquid nitrogen.

### X-ray data collection and structure determination

Cryo-cooled crystals (100 K) were measured using single-wavelength X-ray diffraction experiments at the 19-ID beamline of the Structural Biology Center, Advanced Photon Source at Argonne National Laboratory (we used the SBCcollect program). We integrated, scaled and merged intensities of each data set (HKL-3000 program suite was used^39^). The structure of 3CLpro in complex with masitinib was determined using the molecular replacement method^40^ with an apo form of 3CLpro (PDB code: 7JFQ) as a search template. In the difference Fourier maps, extra electron densities were observed in the substrate binding site of 3CLpro and were subsequently identified as the contribution of masitinib. One data set with a resolution limit up to 1.60 Å was selected for further model rebuilding, including building masitinib into extra densities using the program Coot^41^ and refinement using the program phenix.refine^42^ (Table 1). We used MOLPROBITY^43^ to validate the stereochemistry of the structure (Table 1).

## Synthesis of fluorogenic peptide of 3CLpro kinetics assay

### General

We performed low resolution mass spectral and liquid chromatography analyses using the Advion Expression-L mass spectrometer (Ithaca, NY), which was coupled to an Agilent 1220 Infinity LC System.

### Synthetic protocols

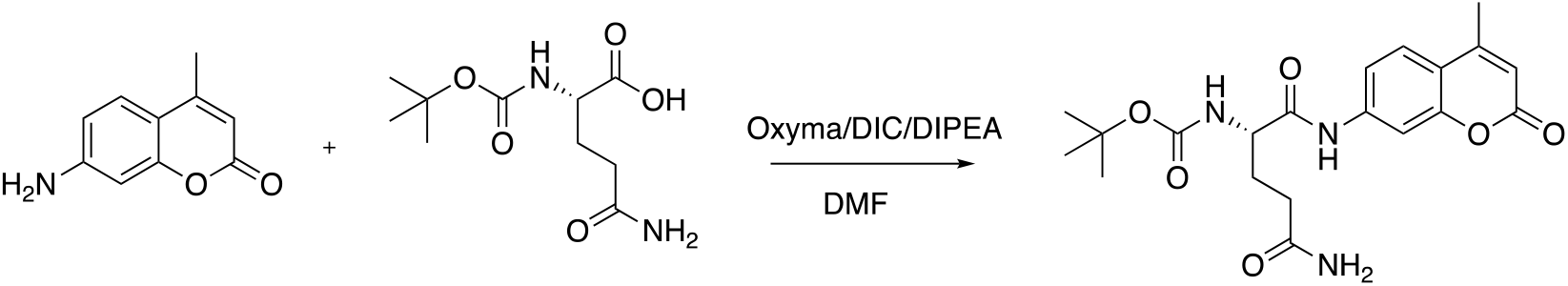

**1**. 7-methyl-4-amino coumarin (AMC) (0.5 g, 2.85 mmol, 1.0 eq) was dissolved in dry DMF (10 mL) in a 25 mL round-bottom flask. Boc-glutamine (1.4 g, 5.7 mmol, 2 eq.) was added, followed by DIPEA (1 mL, 5.7 mmol, 2 eq.), Oxyma (0.811 g, 5.7 mmol, 2 eq.) and DIC (530 μL, 3.4 mmol, 1.2 eq.). The reaction mixture was stirred at room temperature for 4 hours and then heated to 50°C. After stirring overnight, the reaction was cooled and concentrated under reduced pressure. Purification by column chromatography (0-5% methanol in DCM) yielded **1** (0.552 g, 48%). **LRMS-ESI(+):** Calculated for C_20_H_25_N_3_O_6_ [M+H^+^] 403.17, found 403.2

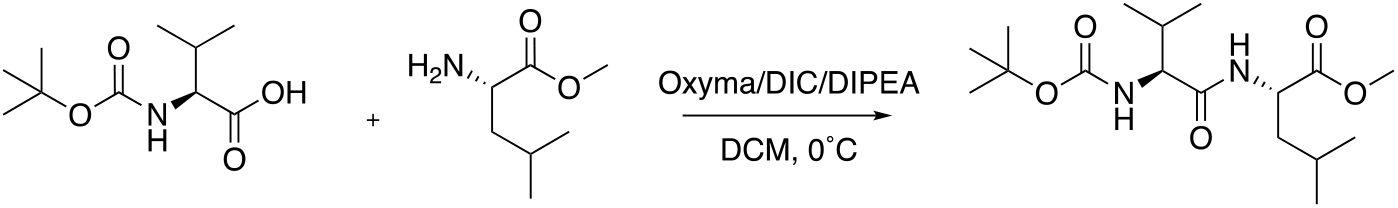

**2**. L-leucine methyl ester (0.30 grams, 1.65 mmol, 1.0 eq.) was dissolved in dichloromethane (10 mL) and, following addition of DIPEA (431 μL, 2.48 mmol, 1.5 eq.) the reaction was cooled to 0°C. In a separate flask, Boc-valine (0.716 g, 3.3 mmol, 2 eq.) and Oxyma (0.352 g, 2.48 mmol, 1.5 eq.) were dissolved in DCM and stirred for 5 minutes, after which DIC (384 μL, 2.48 mmol, 1.5 eq.) was added. This solution was then added to the reaction mixture and stirred for two hours while the solution warmed to room temperature. The solution was then diluted with DCM, washed with brine, and dried over magnesium sulfate. Purification by column chromatography (0-40% EtOAc in hexanes) yielded **2** (0.445 g, 78%). **LRMS-ESI(+):** Calculated for C_17_H_32_N_2_O_5_ [M+H^+^] 344.23, found 344.5.

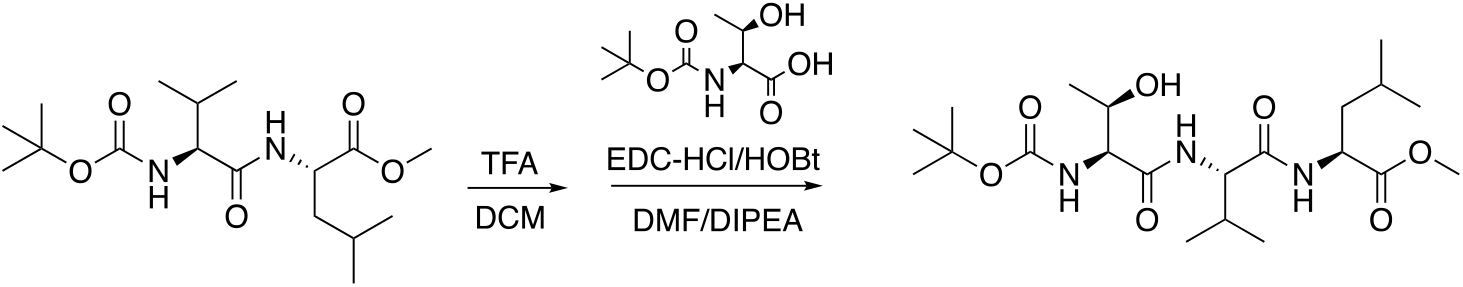

**3**. Compound **2** (0.210 g, 0.610 mmol, 1 eq.) was dissolved in a solution of 25% TFA in DCM and stirred for 30 minutes. After TLC indicated removal of the Boc group, the reaction was concentrated and the product washed with DCM three times. The deprotected peptide was then dissolved in DMF (5 mL) and DIPEA (215 μL, 1.22 mmol, 2.0 eq.), EDC•HCl (0.176 g, 0.914 mmol, 1.5 eq.), and HOBt (0.150, 0.914 mmol, 1.5 eq.) were added. The reaction was stirred for two hours at room temperature and then the solvent was removed under reduced pressure. Purification by column chromatography (0-40% EtOAc in hexanes) yielded **3** (0.445 g, 78%). **LRMS-ESI(+):** Calculated for C_21_H_39_N_3_O_7_ [M+H^+^] 446.46, found 447.

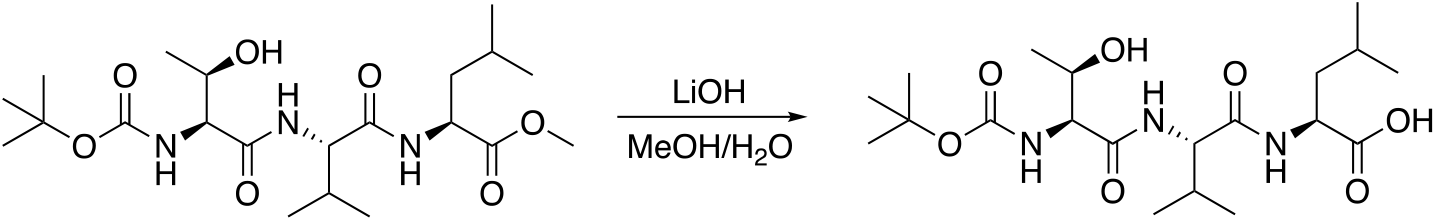

**4**. Compound 3 (0.150 g, 0.336 mmol, 1 eq.) was dissolved in solution of MeOH and water (1:1, 4 mL). Lithium hydroxide (0.016 g, 0.672 mmol, 2 eq.) was added and stirred in suspension. After three hours, reaction as acidified to pH = 2.0 and extracted with ethyl acetate. The organic phase was washed using brine and was dried over magnesium sulfate. Concentration under reduced pressure yield **4** (0.145 g, 98%) as pale yellow oil. **LRMS-ESI(+):**Calculated for C_20_H_37_N_3_O_7_ [M+H^+^] 431.26, found 431.5.

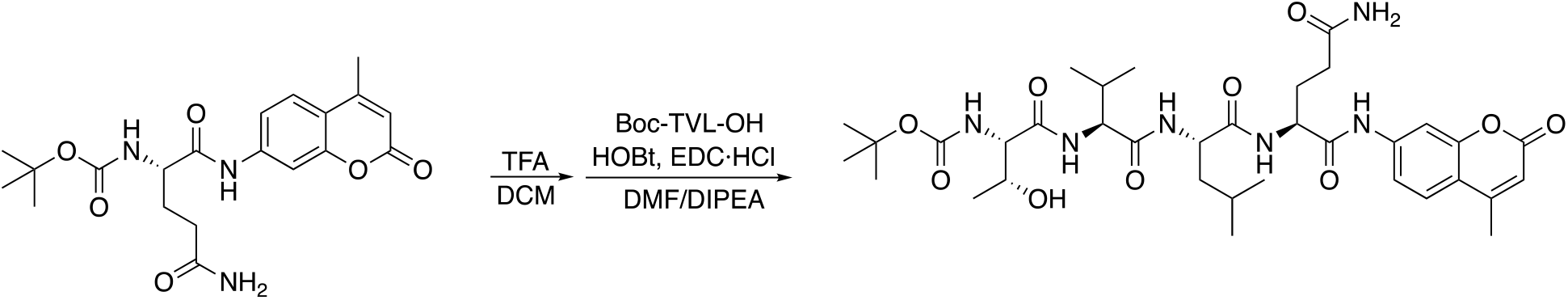

**5**. Compound 1 (0.085 g, 0.211 mmol, 1 eq.) was dissolved in a solution of 25% TFA in DCM and stirred for 30 minutes. After TLC indicated removal of the Boc group, the reaction was concentrated and the product washed with DCM three times. It was then dissolved in DMF (5 mL) and DIPEA (74 μL, 0.42 mmol, 2.0 eq.), EDC•HCl (0.061 g, 0.316 mmol, 1.5 eq.), and HOBt (0.034, 0.211 mmol, 1.0 eq.) were added. The reaction was stirred for two hours at room temperature and then the solvent was removed under reduced pressure. Purification by column chromatography (1 to 10% MeOH in DCM) yielded **5** (0.085 g, 56%). **LRMS-ESI(+):** Calculated for C_35_H_52_N_6_O_10_ [M+H^+^] 716.37, found 716.4.

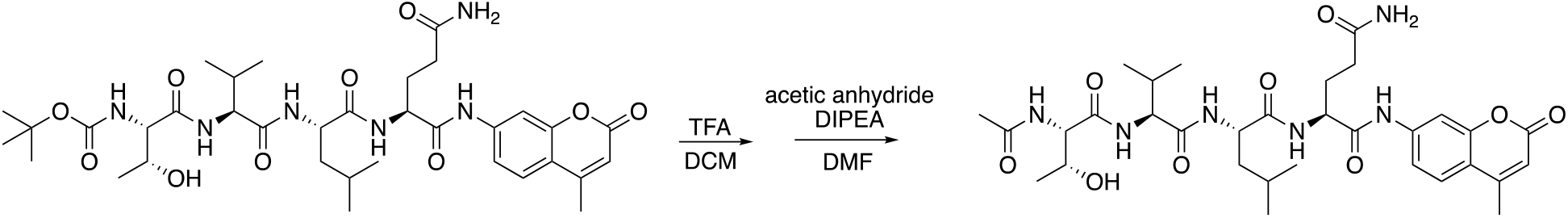

**6**. Compound 5 (0.055 g, 0.0893 mmol, 1 eq.) was dissolved in a solution of 25% TFA in DCM and stirred for 30 minutes. After TLC indicated removal of the Boc group, the reaction was concentrated and the product washed with DCM three times. It was then dissolved in DMF (2 mL) with DIPEA (32 μL, 0.179 mmol, 2 eq.), and acetic anhydride (18 μL, 0.179 mmol, 2 eq.) was added slowly. After stirring for 10 minutes, solvent was removed under reduced pressure. The crude reaction mixture was re-dissolved in water and washed with dichloromethane. The aqueous layer was then cooled to 0°C, upon which action **6** precipitated as a white solid (**LRMS-ESI(+):** Calculated for C_32_H_36_N_6_O_9_ [M+H^+^] 658.33, found 658.5.

### Picornaviruses infection

Prior to infection, Huh7 cells were pretreated for two hours. Virus was diluted using serum-free DMEM (SFM) to achieve an MOI of 0.01. Cell supernatants (collected at 24 hrs post infection) were dilutions in SFM and used to inoculate Vero cells for 10-15 min at 37⁰C. Cells were incubated for 2 days at 37⁰C after overlaying them DMEM containing 2% NBCS and 0.8% agarose. Cells were then fixed with 4% formalin and revealed with crystal violet solution (10% crystal violet; Sigma-Aldrich). The number of plaque forming units (PFU/per milliliter) were then calculated.

### 3C protease activity assay

Huh7 cells were transfected with LipoD293 (SignaGen Laboratories) with 3C substrate, 3C protease (derived from CVV3) and a Renilla transfection control plasmid (siCheck). Protease and target constructs were generated using protocols previously described^28^. The cells were combined with firefly substrate (Bright-Glo; Promega) followed by subsequent Renilla (Stop and Glo; Promega) luciferase substrate 24 hours post transfection. Assays were performed using the manufacturer’s recommendations (Promega) and a Veritas Microplate Luminometer (Turner BioSystems) was used to quantify the results.

### Influenza A infection

MDCK-SIAT1-TMPRSS2 cells were infected with Influenza A/Puerto Rico/8/1934 (PR8) at an MOI of 0.01 TCID50/cell. Following a 1 hour adsorption, virus was removed and the cells were washed. Viral growth medium was added with either masitinib or DMSO to a final concentration of 10μM. We harvested and clarified supernatants at 20 hours post infection. We titrated the supernatants using TCID on MDCK-SIAT1-TMPRSS2 cells.

### LCMV infection

One day prior to infection, A549 cells were seeded in 12 well dishes (80,000 cells per well). Cells were infected with rLCMV at an MOI of 0.01 for one hour at 37°C. The inoculum was removed and cells were overlaid with 1 ml of complete media containing masitinib or DMSO only control. Supernatants were harvested at 48 hours after infection, were clarified and titrated by a previously described immuno-focus assay^44,45^, using a mouse anti-LCMV nucleoprotein antibody (1-1.3, kindly provided by M. Buchmeier) and a peroxidase-labeled goat anti-mouse antibody (SeraCare).

### Measles Virus Inhibition assay

Vero cells were infected with a luciferase-expressing Measles virus at an MOI of 0.01 for 90 minutes. We removed the inoculum and added fresh medium containing masitinib or DMSO to the cells for further culture. Three days later, firefly luciferase activity was measured by adding 0.5mM of D-Luciferin to each well and was quantified with an Infinite M200 Pro multimode microplate reader.

### Statistical analyses

For all experiment described, the size of the sample (n) refers to independent biological samples tested. All analyses were performed in Matlab. Multiple-comparison corrections was performed using the FDR method.

### Data Availability

All data are available from the corresponding author upon reasonable request. The X-ray structure of 3CLpro-bound masitinib has been deposited to PDF under accession number 7JU7.

---

**Figure S1.**
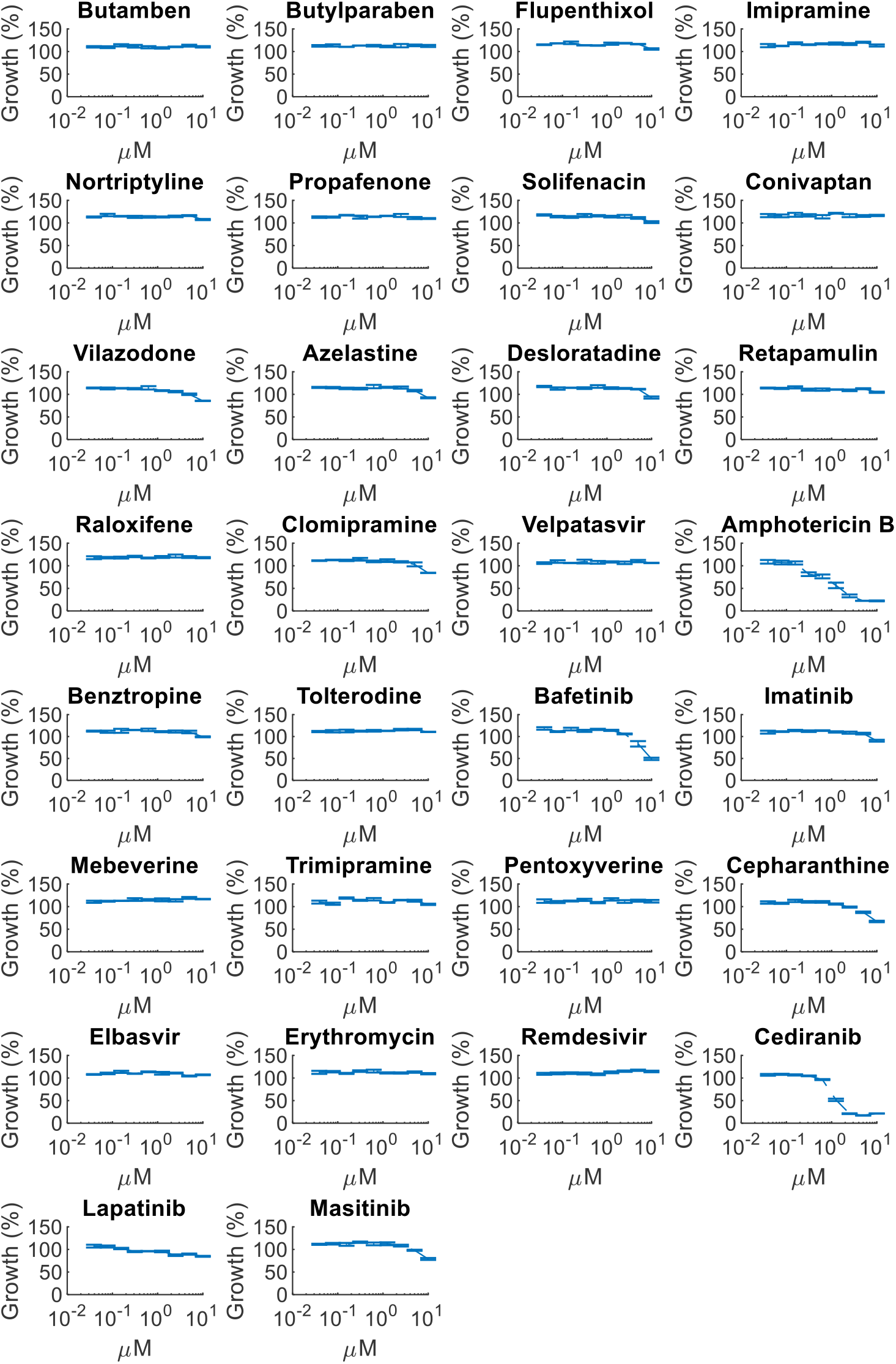
Effect of identified drugs on cellular growth. Blue lines show the effect of the drugs on cell growth (% of no drug controls). Mean±S.E. n=3.

**Figure S2.**
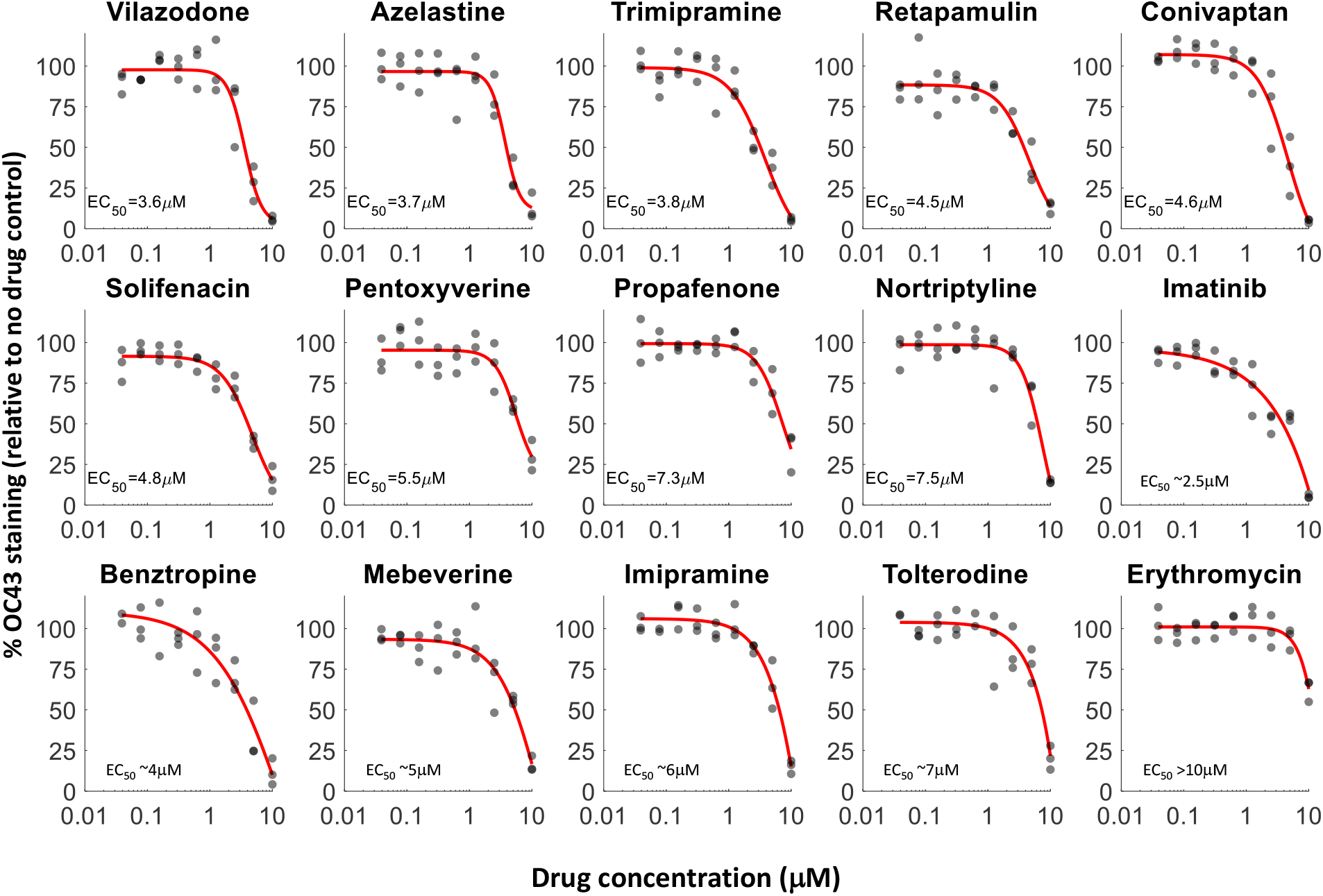
Additional dose-response curves for OC43 infection. Dose response curves of the remaining tested drugs, n = 3. Individual measurements are shown as semi-transparent circles (note that some circles overlap).

**Figure S3.**
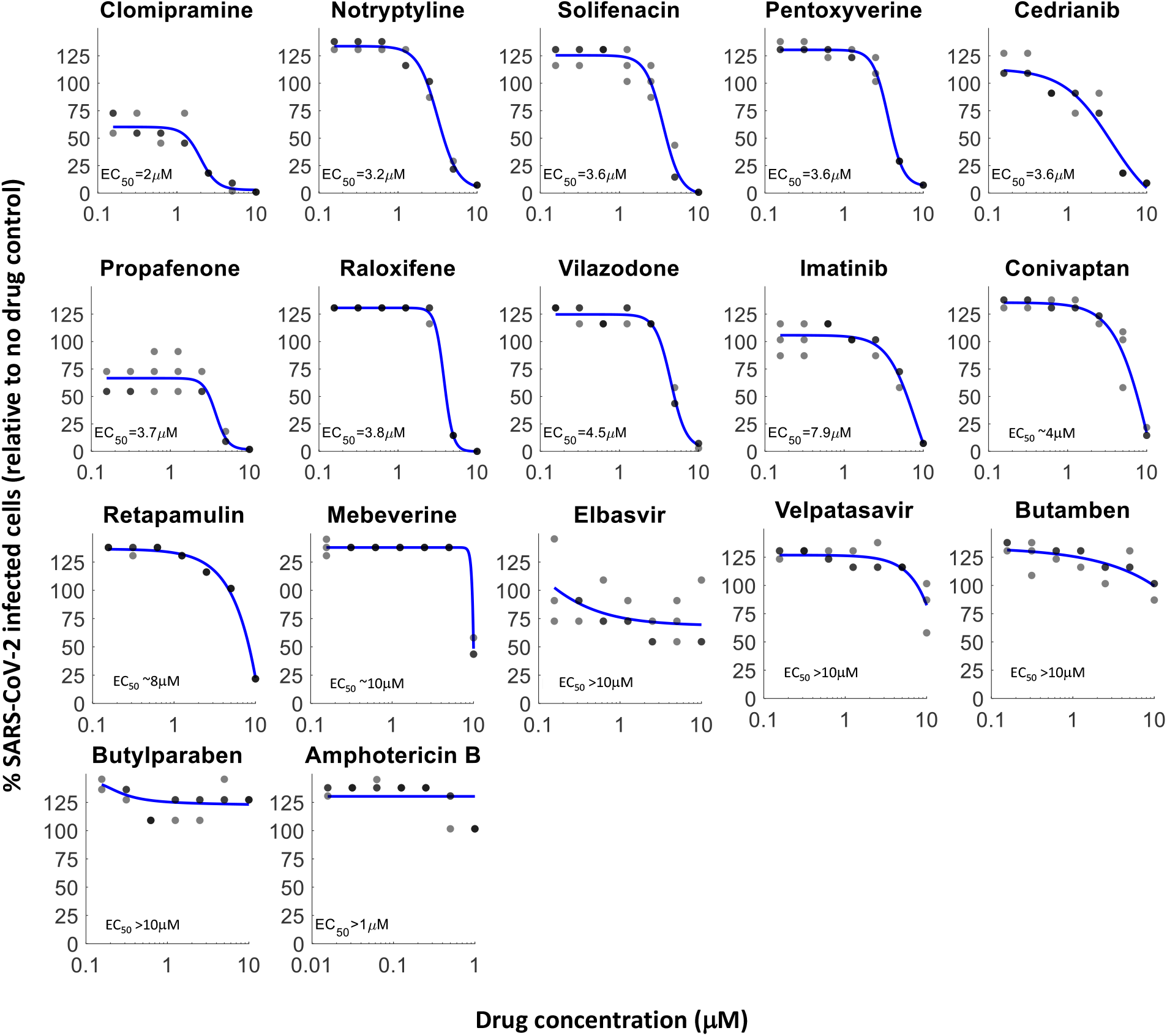
Additional dose-response curves for SARS-CoV-2 infection. Dose response curves of the remaining tested drugs, n = 3. Individual measurements are shown as semi-transparent circles (note that some circles overlap).

**Figure S4.**
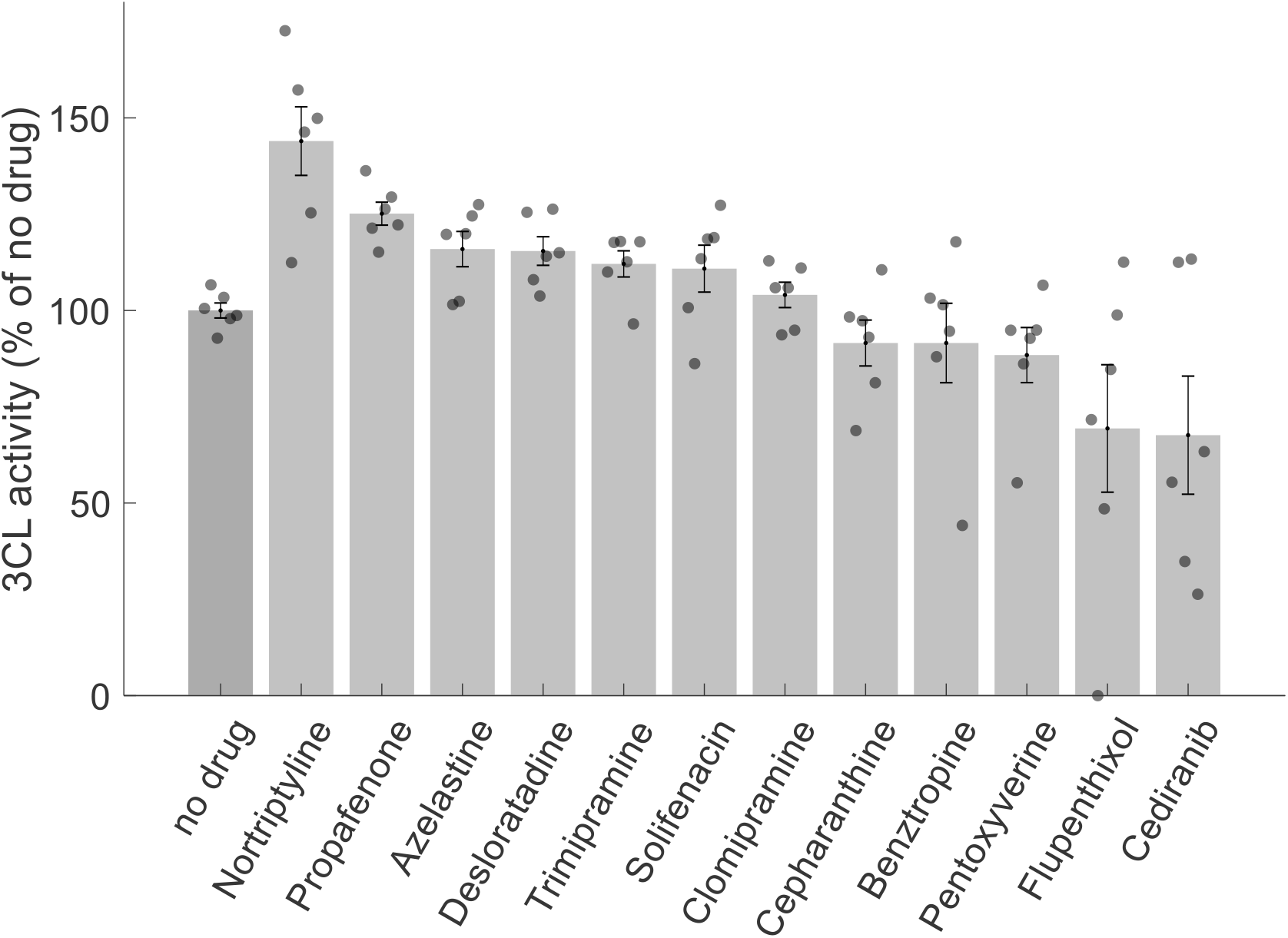
Effect of remaining tested drugs on 3CLpro FlipGFP reporter assay. A FlipGFP reporter assay was performed to screen for potential inhibition of 3CLpro by the identified drugs at a single concentration (10μM). Shown are all the drugs that did not show a significant reduction in 3CLpro activity (p-value>0.05, one-tailed t-test, FDR-corrected). n = 6. Individual measurements are shown in circles. Bars depict mean ± s.e. The drugs that showed statistically significant inhibition of 3CLpro are shown in Figure 3A.

**Figure S5.**
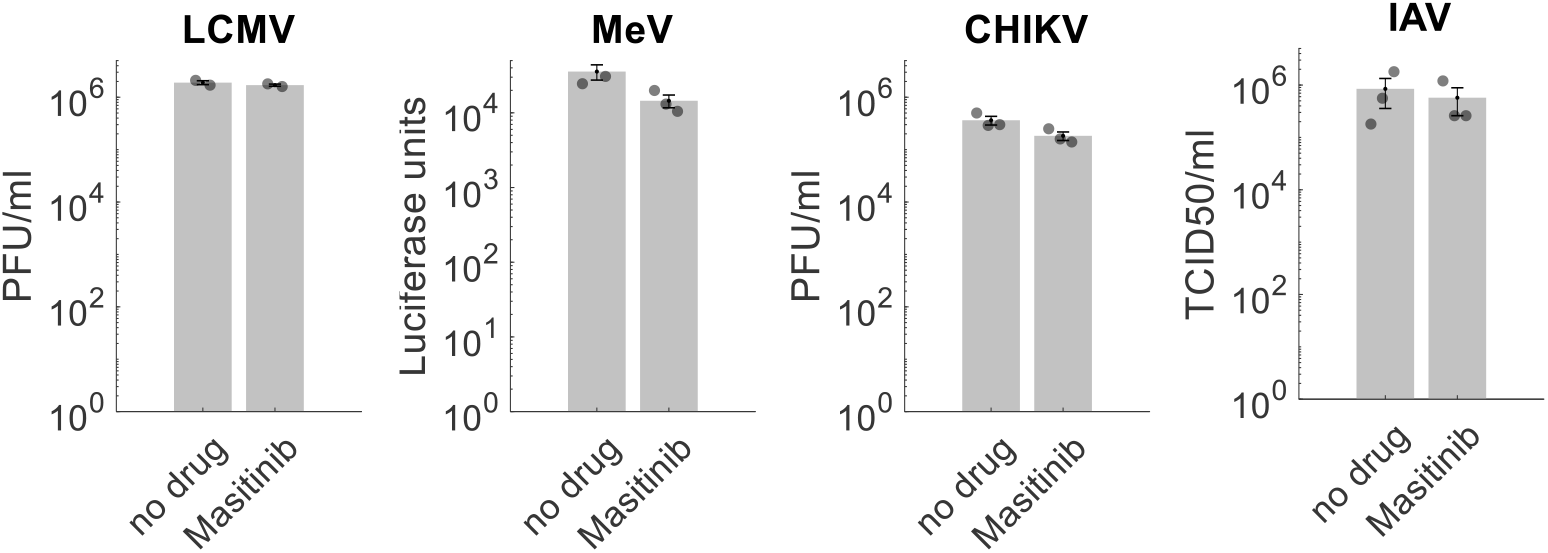
Masitinib does not inhibit other RNA viruses. Masitinib (10μM) did not show a significant effect on cells infected by influenza A virus (IAV, Orthomyxoviridae), measles virus (MeV, Paramyxoviridae), lymphocytic choriomeningitis virus (LCMV) and Chikungunya virus (CHIKV, Togaviridae). n=3 for all except LCMV (n=2). p-values>0.07 (one-tailed t-test, FDR-corrected).

