## Supplementary material for "Drug repurposing screen identifies masitinib as a 3CLpro inhibitor that blocks replication of SARS-CoV-2 *in vitro*": Table_S2_EC50_OC43_SARS_CoV_2

| # | Drug | EC50 OC43 | EC50 SARS2 | Structure |
| --- | --- | --- | --- | --- |
| 1 | Butamben (n-butyl-4-aminobenzoate) | 3.1 | >10 | 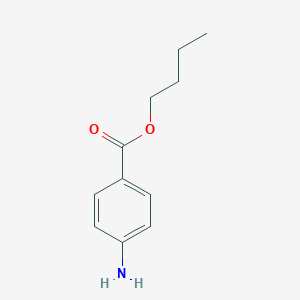 |
| 2 | Butylparaben (n-buytl-4-hyfroxybenzoate) | 1.7 | >10 | 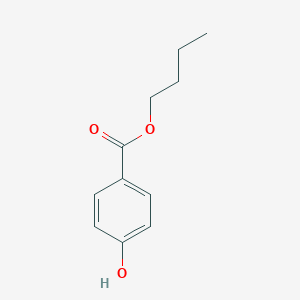 |
| 3 | Flupenthixol dihydrochloride | 1.8 | 0.56 | 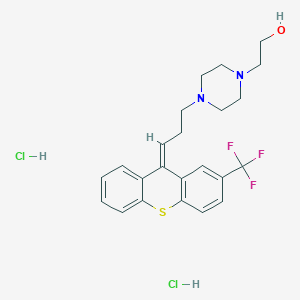 |
| 4 | Imipramine HCl | 6 | ND | 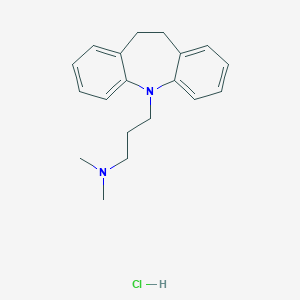 |
| 5 | Nortriptyline hydrochloride | 7.5 | 3.2 | 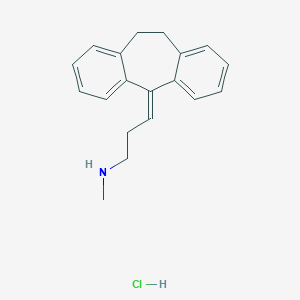 |
| 6 | Propafenone HCl | 7.3 | 3.7 | 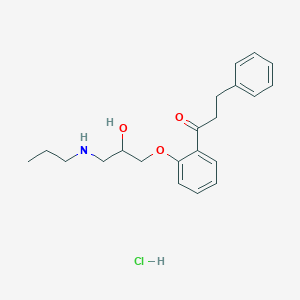 |
| 7 | Solifenacin succinate | 4.8 | 3.6 | 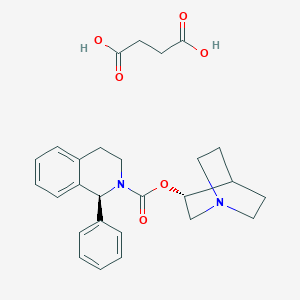 |
| 8 | Conivaptan HCl | 4.6 | 4 | 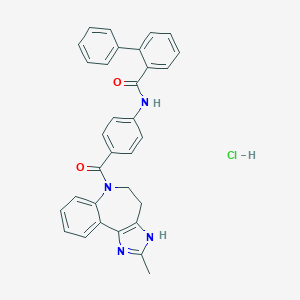 |
| 9 | Vilazodone HCl | 3.6 | 4.5 | 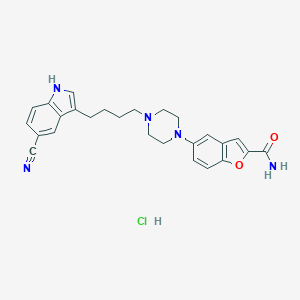 |
| 10 | Azelastine HCl | 3.7 | 2.4 | 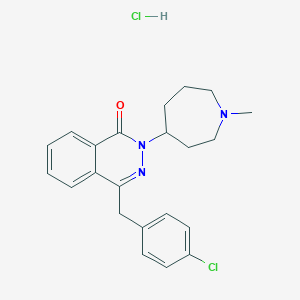 |
| 11 | Desloratadine | 3.2 | 0.9 | 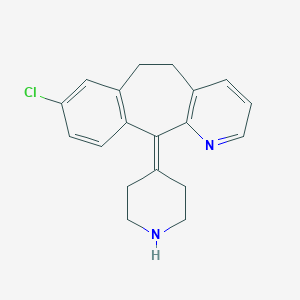 |
| 12 | Retapamulin | 4.5 | 8 | 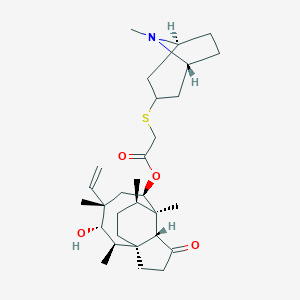 |
| 13 | Raloxifene HCl | 2.7 | 3.8 | 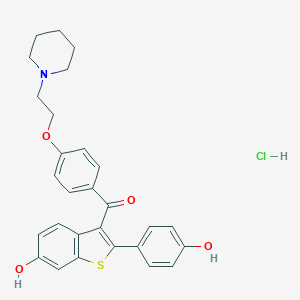 |
| 14 | Clomipramine HCl | 3 | 2 | 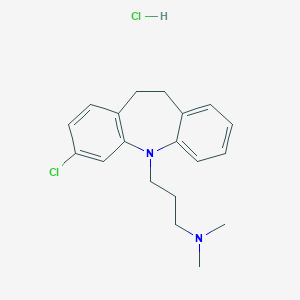 |
| 15 | Velpatasvir | 2 | >10 | 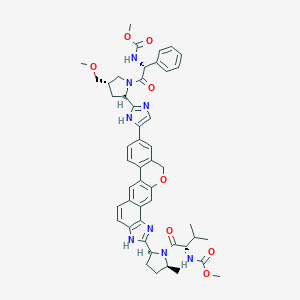 |
| 16 | Amphotericin B | 0.04 | >1 | 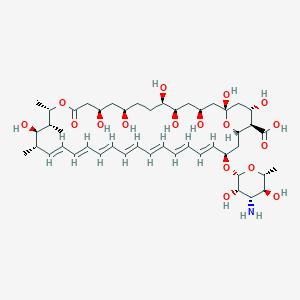 |
| 17 | Benztropine mesylate | 4 | 1.8 | 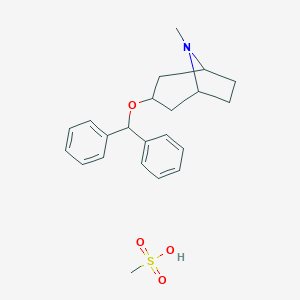 |
| 18 | Tolterodine tartrate | 7 | ND | 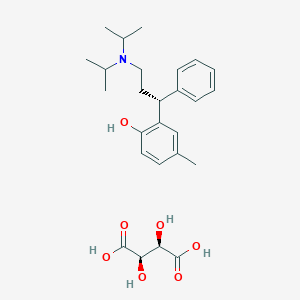 |
| 19 | Bafetinib (INNO-406) | 2.1 | 2.2 | 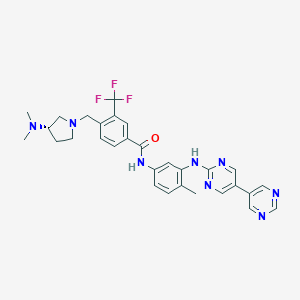 |
| 20 | Imatinib (STI571) | 2.5 | 7.9 | 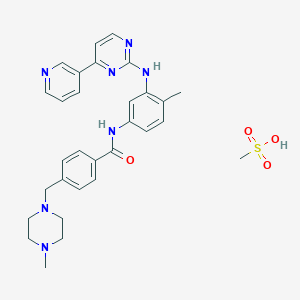 |
| 21 | Mebeverine Hydrochloride | 5 | 10 | 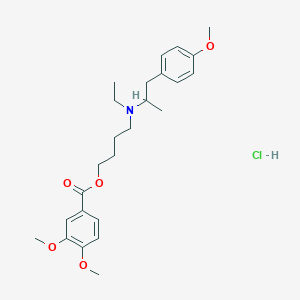 |
| 22 | Trimipramine maleate | 3.8 | 1.5 | 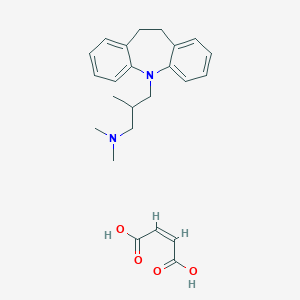 |
| 23 | Pentoxyverine Citrate | 5.5 | 3.6 | 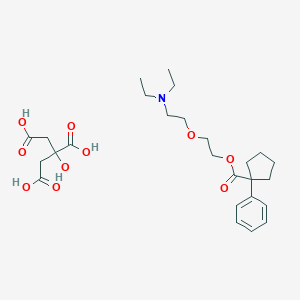 |
| 24 | Cepharanthine | 0.77 | 0.13 | 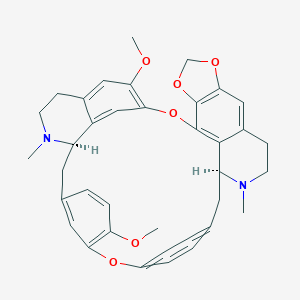 |
| 25 | Elbasvir | 0.17 | >10 | 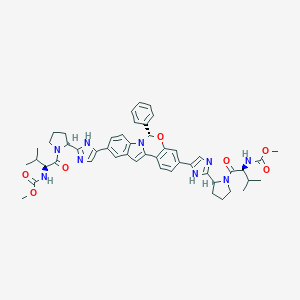 |
| 26 | Erythromycin Cyclocarbonate | >10 | ND |  |
| 27 | Remdesivir | 0.96 | 0.1 |  |
| 28 | Cediranib (AZD2171) | 0.5 | 3.6 | 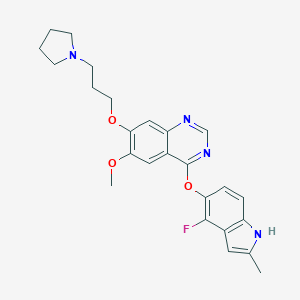 |
| 29 | Lapatinib | 3.1 | 1.6 | 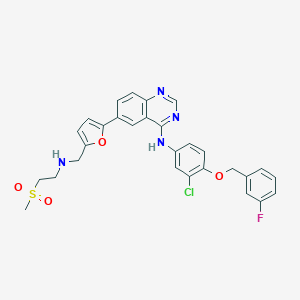 |
| 30 | Masitinib (AB1010) | 2.1 | 3.2 | 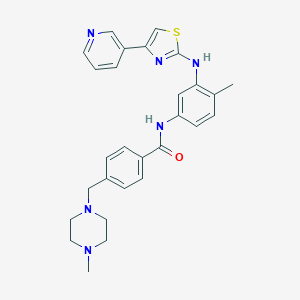 |
